# Consistent and reproducible long-term *in vitro* growth of health and disease-associated oral subgingival biofilms

**DOI:** 10.1101/324475

**Authors:** Irina M. Velsko, Luciana M. Shaddox

## Abstract

**Background**: Several *in vitro* oral biofilm growth systems can reliably construct oral microbiome communities in culture, yet their stability and reproducibility through time has not been well characterized. Long-term *in vitro* growth of natural biofilms would enable use of these biofilms in both *in vitro* and *in vivo* studies that require complex microbial communities with minimal variation over a period of time. Understanding biofilm community dynamics in continuous culture, and whether they maintain distinct signatures of health and disease, is necessary to determine the reliability and applicability of such models to broader studies. To this end, we performed next-generation sequencing on biofilms grown from healthy and disease-site subgingival plaque for 80 days to assess stability and reliability of continuous oral biofilm growth.

**Results**: Biofilms were grown from subgingival plaque collected from periodontitis-affected sites and healthy individuals for ten eight-day long generations, using hydroxyapatite disks. The bacterial community in each generation was determined using Human Oral Microbe Identification by Next-Generation Sequencing (HOMINGS) technology, and analyzed in QIIME. Profiles were steady through the ten generations, as determined by species abundance and prevalence, Spearman’s correlation coefficient, and Faith’s phylogenetic distance, with slight variation predominantly in low abundance species. Community profiles were distinct between healthy and disease site-derived biofilms as demonstrated by weighted UniFrac distance throughout the ten generations. Differentially abundant species between healthy and disease site-derived biofilms were consistent throughout the generations.

**Conclusions**: Healthy and disease site-derived biofilms can reliably maintain consistent communities through ten generations of *in vitro* growth. These communities maintain signatures of health and disease and of individual donors despite culture in identical environments. This subgingival oral biofilm growth and perpetuation model may prove useful to studies involving oral infection or cell stimulation, or those measuring microbial interactions, which require the same biofilms over a period of time.

## Background

The oral biofilm is a complex microbial community that is responsible for the world’s most common diseases, periodontal disease [1] and caries [2]. Ecological theories of oral infectious diseases such as Marsh’s ecological plaque hypothesis [3] and the recent polymicrobial synergy and dysbiosis model of disease development [4] emphasize the role of complex community changes as drivers of disease. It is now clear that oral biofilms differ between sites of health and disease at the gene/genomic level [5], the transcriptional level [6,7], and the metabolic level [8]. However, despite substantial research into cause of these infectious oral diseases, relatively little progress has been made in understanding how community dynamics and species interactions drive disease. Studies employing polymicrobial communities, which are necessary to elucidate oral disease pathology, are rare due to the difficulty of independently culturing many oral bacterial species, and the uncertainty in working with undefined samples derived from dental plaque or saliva.

Differences in immunological and physiological responses to stimulation with bacterial structures such as LPS, single bacterial species, and multiple bacterial species *in vitro* and *in vivo* have demonstrated that it is necessary to study polymicrobial-induced diseases with complex communities to better understand disease etiology. For example, murine oral co-infection with *Tannerella forsythia* and *Fusobacterium nucleatum* [9] or with *Porphyromonas gingivalis* and *Treponema denticola* [10] induces greater alveolar bone resorption than either species alone, while a polybacterial inoculum of *P. gingivalis*, *T. denticola, T. forsythia*, and *F. nucleatum* elicits a unique immune/inflammatory response and physiologic changes compared to individual oral infection with each of the species [11–16]. Additionally, peripheral blood mononuclear cells (PBMCs) from localized aggressive periodontitis (LAP) patients and healthy patients respond uniquely to stimulation with bacterial surface components, intact complex oral biofilms, and dispersed complex oral biofilms [17] indicating that the choice of stimulant in immunological assays can skew the conclusions drawn about cellular responses in infectious diseases. In contrast, cytokine responses of whole blood from both LAP and periodontally healthy individuals are indistinguishable between stimulation with LAP site-derived and healthy site-derived dental plaque [17]. Additionally, unique subgingival plaque community profiles correlate with distinct gingival crevicular fluid cytokine profiles in generalized aggressive periodontitis patients and healthy patients [18], chronic periodontitis patients, and clinically healthy sites in chronic periodontitis patients [19].

Bacterial-bacterial interactions in biofilms, particularly through metabolite crossfeeding, influence bacterial physiology, which in turn directs host immunological and physiological responses to the biofilm. Bacteria in dental plaque have complex nutritional relationships that determine community composition and structure [20,21], by enhancing and promoting growth of neighboring species [22,23], and altering expression of surface markers, such as LPS [24,25]. Comparative genome analysis of *P. gingivalis, T. denticola*, and *T. forsythia* revealed that certain metabolic pathways are covered between the three that are not complete in any individually [26], suggesting that close proximity of these species may be necessary for specific metabolite use. Such intimate dependency may explain their frequent co-detection in plaque samples, which led to their grouping as Socransky’s “red complex”, particularly and strongly associated with periodontal disease [27].

Several reliable, reproducible *in vitro* complex biofilm models for studying oral biofilm microbe-microbe interaction and biofilm-host interaction exist, based on saliva and dental plaque inocula [28–32]. The subgingival plaque growth model developed by Walker and Sedlacek [28] demonstrated, using DNA-DNA checkerboard hybridization and scanning electron microscopy, that *in vitro* biofilm growth and development closely resembled the known succession of bacteria in dental plaque, and the mature community at ten days growth had species composition and proportions highly similar to the original plaque inocula [28]. There were promisingly also distinguishable differences between biofilms grown from healthy site plaque and periodontitis-affected site plaque. Further characterization of this model, again by culture-dependent DNA-DNA checkerboard hybridization, found it possible to perpetuate the biofilms up to seven times/generations and maintain the community composition [33]. Here we have further characterized this model by using Next Generation Sequencing (NGS) to trace the community dynamics of subgingival biofilms grown from healthy site and periodontitis-affected site plaque through ten generations of subcultures. We demonstrate stable biofilm communities that maintain differences between healthy site and periodontitis-affected site plaque inocula, as well as between individual donors, making it a useful model for long-term study of biofilm microbe-microbe interactions and biofilm-host interactions.

## Materials and Methods

### Collection of subgingival biofilms and saliva

Subgingival plaque samples were collected as described in [35]. In brief, four subgingival biofilm samples were collected by paper point for growth and propagation: one from each of two periodontally healthy subjects, and one from a localized aggressive periodontal disease (LAP)-affected site in each of two LAP patients. Samples were collected as part of a larger clinical study approved by the University of Florida Institutional Review Board (IRB #201400349, ClinicalTrials.gov NCT01330719). The LAP patients had at least 2 sites with ≥5mm pocket depth and a history of site progression based on radiographic assessment, attachment loss of ≥3mm and bone loss on first molar or incisor and in no more than two teeth affected by bone loss other than first molar and incisor, and had not yet received any periodontal treatment in the previous 6 months, while periodontally healthy individuals presented with no pocket depth ≥5mm and absence of clinical attachment loss by full mouth probing and no evidence of bone loss by radiographic imaging [34,35]. Each plaque sample was collected from a first molar. Paper points were immediately placed in tubes containing reduced Ringer’s solution and stored at 4°C while saliva was collected and prepared. From the same four patients, 5 mL unstimulated saliva was collected in a 50mL tube and diluted 1:10 in reduced Ringer’s solution [33]. Diluted saliva samples were centrifuged at 1200 g × 10 min to remove large particulate matter then filter-sterilized through a 0.2μm membrane, divided into 2 mL aliquots and stored at −20°C.

### Initiation of biofilm growth from subgingival plaque samples

Biofilms were grown as previously described [33,35] (Fig. 1). In brief, subgingival plaque samples were gently sonicated in a water bath for 30 seconds to break up the biofilm and stored at 4°C while hydroxyapatite (HA) disks were prepared for biofilm inoculation. Hydroxyapatite disks were placed into wells of two sterile 24-well cell culture plates, one for disease-site biofilms and one for healthy-site biofilms, with two wells for each biofilm source to ensure growth (four biofilm lineages). The HA disks were coated for 2 hours at room temperature with sterile 1:10 saliva from the donor whose plaque sample was later added to the well. The saliva was removed and 2 mL sterile tryptic soy broth supplemented with 5μg/mL hemin and 1μg/mL menadione (TSB-hk) gently added to each well. Plaque samples were resuspended by briefly vortexing, and wells were inoculated with 50 μL of plaque sample from the same donor as the saliva used to coat the wells. Plates were placed in an anaerobic incubator at 37°C-75%N_2_/10%CO_2_/10%H_2_ for static growth. Biofilms were grown in replicates for immune cell stimulation experiments for a separate study [35].

**Fig. 1.**
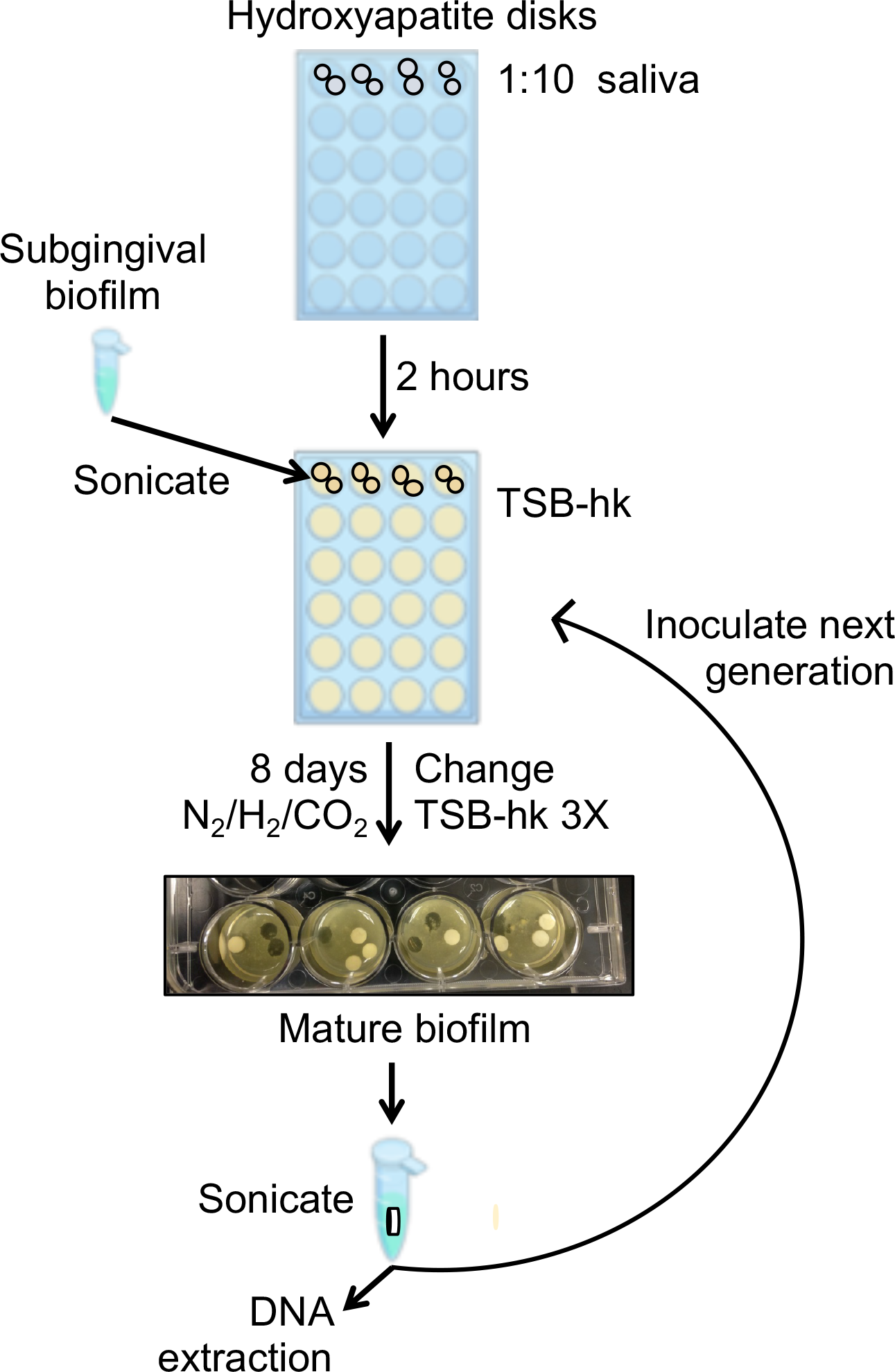
Schematic diagram the biofilm inoculation and propagation. Each biofilm lineage was propagated in an independent 24-well plate. For details see the methods section.

### Propagation of subgingival biofilms *in vitro*

Biofilms were grown for 8 days before dispersal and re-inoculation (Fig. 1), as described in [35]. Every 48 hours growth media was gently removed from wells and replaced by slowly adding a fresh 2 mL of reduced TSB-hk. On the 8^th^ day, new HA disks were added to new wells and coated with respective 1:10 sterile saliva for 2 hours in the anaerobic chamber, then saliva was removed and replaced with 2 mL fresh reduced TSB-hk. A single disk with mature biofilm (8^th^ day of growth) from each of the 4 sources was removed from a well and placed in a tube with 1 mL reduced Ringer’s solution. The tubes were sonicated in a water bath for 30 sec, vortexed briefly to disperse any remaining clumps, and 50 μL of the suspension used to inoculate the well for the second generation. This procedure was repeated for 10 generations (80 days) for all 4 biofilm sources. The biofilm suspension was further used for viable cell count estimations and DNA extraction.

### Viable cell count estimations

The biofilm suspension was additionally plated to estimate viable cell counts on the HA disk at the time of dispersal. For each dispersed biofilm sample, serial dilutions of 100 μL from 10^−1^ – 10^−7^ were made in sterile reduced Ringer’s solution, and 100 μL of dilutions 10^−4^ – 10^−7^ were plated in duplicate on reduced TSB-hk blood agar plates. Plates were grown for 5 days in the anaerobic chamber, then colonies counted and plate counts averaged for viable cell count estimations.

### DNA extraction and HOMINGS 16S rRNA gene sequencing

The remaining biofilm suspension from each generation 1-10 was spun at 10000 rpm for 2 min to pellet cells, the supernatant was removed and the pellet stored at −20°C for DNA extraction. DNA was extracted from the pellet of each biofilm source for all 10 generations as well as a plaque sample from one of the two LAP patient plaque donors with a QIAGEN kit, and sent to the Forsyth Institute for next generation sequencing with Human Oral Microbe Identification using Next Generation Sequencing (HOMINGS) technology [36]. In brief, HOMINGS uses V3-V4 region-specific 16S rRNA gene primers for amplification and library generation, then libraries are sequenced on an Illumina MiSeq. Reads are identified using the program ProbeSeq by matching to primers that uniquely identify the 16S rRNA gene of 638 bacterial and archaeal species known to reside in the oral cavity based on the Human Oral Microbiome Database. Reads not matching species probes are assigned to a genus when possible, and all remaining reads are unmatched. Raw read data was processed by the Forsyth Institute/HOMINGS and a table with read counts and taxa counts was generated. All analyses were performed on the species absolute read count table generated by HOMINGS analysis(Table S1).

A separate identification procedure was performed in QIIME to confirm the trends seen with HOMINGS technology. Reads were joined and demultiplexed in QIIME v1.9 [37], using a quality score cut-off of q ≥ 30, using scripts join_paired_ends.py and split_libraries_fastq.py, respectively. Read quality was checked using FastQC [38]. Taxonomic identification was performed using pick_closed_reference_otus.py in default (uclust) with the additional flag enable_rev_strand_match, using the GreenGenes 13_8 reference database with clusters set at 97% identity. Operational taxonomic unit tables were generated in QIIME with summarize_taxa.py at the genus (L6) and species (L7) levels. The OTU tables are presented in Table S2.

### Community diversity analyses

QIIME v1.9 was used for all community diversity analyses [37]. Several species have multiple probes in HOMINGS and share the same Human Oral Taxon (HOT) number, and counts for these species were summed to remove duplicate HOT numbers for all analyses. The species tables within each of the 4 biofilm lineages were compared for consistency through generations by Spearman correlation in QIIME. Generations were tested one to the next as 1-2, 2-3, 3-4, 4-5, 5-6, 6-7, 7-8, 8-9, and 9-10, as well as 1-10, in order to correlate to the original generation. Diversity within the different biofilms from LAP disease sites and within healthy sites (alpha diversity) was assessed using the metrics Faith’s phylogenetic distance, Chao1, observed OTUs and Shannon entropy, on tables rarefied to a depth of 20,000 reads/sample. Rarefications were performed and alpha diversity metrics calculated 10 times and then averaged. Diversity between biofilms from LAP disease and healthy sites (beta diversity) were calculated using weighted UniFrac distance [39,40] and the Sorensen-Dice index. Principal coordinates analysis was performed on beta diversity matrices from both metrics and the results were plotted in QIIME. Jack-knifed beta diversity was also determined by rarefying the table to 20,000 reads/sample 10 times and calculating diversity metrics. Principal coordinate analysis was likewise performed on both beta diversity matrices and the results were plotted in QIIME, while UPGMA trees based on the beta diversity matrices were generated in QIIME. The core microbiome of health-and disease-site-derived biofilms was calculated in QIIME as species present in all 10 generations of heathy and disease site-derived lineages.

### Differential species analyses

The program Statistical Analysis of Metagenomic Profiles (STAMP) [41,42] was used to assess significantly different species between health and disease overall (two-group analysis) as well as within each generation (two-sample analysis), by comparing the difference in proportions of species between health-and LAP-derived biofilms. For comparison by two-sample analyses, the read counts for each species were summed for both healthy site-derived biofilms and for both LAP site-derived biofilms. All input tables were unrarefied absolute read counts.

### Statistical analyses

Statistical significance of the difference in alpha diversity metrics between healthy-and disease-site derived biofilms were determined in QIIME using a nonparametric Monte Carlo permutations test. Significance of the difference in beta diversity metrics between healthy-and disease-site derived biofilms were determined in QIIME using adonis from the vegan R package. Alpha values of ≤ 0.05 were considered significant in all analyses. Two group analyses in STAMP used White’s non-parametric t-test with bootstrapping to determine the difference between proportion (DP) with cut-off 95% and Storey’s FDR correction, and corrected p-values (q-values) of ≤ 0.05 together with an effect size > 2 were considered significant. Two sample analyses in STAMP used a two-sided Fisher’s exact test with Newcomb-Wilson DP (cut-off 95%) and Storey’s FDR. Corrected p-values (q-values) of ≤ 0.05 and an effect size > 2 were included for comparison in all STAMP analyses.

## Results

### Characteristics of biofilm growth

Biofilm growth was visibly apparent in all inoculated wells after 48 hours of growth, covering the HA disks, the bottom and the sides of the well. By eight days of growth the biofilms covering the HA disks were thick and textured (Fig. 1), with no observable difference between any of the four lineages. When removed from the anaerobic chamber for media changing and propagation, the biofilms grown from disease site plaque smelled distinctly different from the biofilms grown from healthy site plaque, indicating production of more unpleasant volatile compounds, a known characteristic of disease-associated oral biofilms. Viable cell counts remained consistent for each lineage for each generation, reaching ~3×10^8^ cells on the disk. Colonies growing on blood agar plates from all four lineages were varied in size, color and morphology, indicating a polymicrobial community. Disease-derived biofilm samples grew small black-pigmented colonies on blood agar up to three generations, an observed characteristic of disease site dental plaque samples, while both health and disease-derived samples grew brown pigmented colonies for all ten generations, indicating heme-accumulating organisms in both healthy and disease site plaque-derived biofilms.

### Community consistency through generations based on HOMINGS sequencing

Samples had between 50,000 and 164,000 total reads, from which 25%-59% of reads identified at the species level and 6%-40% identified at the genus level (Table S1). The total number of species identified in 3 of the biofilm lineages decreased throughout the ten generations, HB1 from 60 to 56, DB2 from 116 to 64, DB3 from 83 to 52, while the number of species in HB2 increased from 56 to 75 (Table S1) but remained in the range reported for oral biofilms [43]. The healthy site-derived biofilms averaged 59 species each across the 10 generations, while DB2 averaged 83 species and DB3 averaged 67 across the 10 generations. The DB3 plaque inoculum had 154 species detected, and the observed reduction of species detected in biofilms derived from this plaque is consistent with other models of oral biofilm *in vitro* growth [28–32]. Our model supports 53-80% of the initial species based on comparison of the plaque inocula for DB3 and the biofilms of that lineage at each generation. The number of species-identified reads in each lineage however decreased from the first to the tenth generation, ie in HB1 from 76,094 to 40,927, HB2 from 66,206 to 31,677, DB2 from 49,189 to 20,239, and DB3 from 28,597 to 28,122. However, some species that disappear in a specific generation were again detected in subsequent remaining generations. The disappearing-one-generation-reappearing-the-next pattern seen for many of the species detected suggests that they are present through the entire study, and are simply not detected, which may be a result of low abundance in the biofilm, DNA extraction and amplification bias, different sequencing depths, or sequencing errors.

Our independent QIIME analysis classified a comparable number of reads as HOMINGS (3226031 QIIME, 2383617 HOMINGS), and QIIME had nearly half the number of unmatched reads (808149 QIIME, 1869947 HOMINGS) (Table S2). However, QIIME taxonomic assignment had lower resolution, identifying approximately half the number of species in each biofilm, demonstrating the superior taxonomic assignment provided by the species-specific probes used in HOMINGS. The taxonomic profile identified by QIIME was similar to that of HOMINGS, with the majority of assignments belonging to genera and species known to inhabit the oral biofilm.

### Community diversity within healthy and diseased site-derived biofilm lineages

There are conflicting reports on the differences in microbial diversity at healthy tooth sites and those experiencing periodontal disease [5,44,45]. Therefore, we calculated community diversity scores to assess differences between healthy-and LAP site-derived biofilms using Faith’s phylogenetic distance, and observed species. Rarefaction curves show slightly higher diversity in LAP site-derived biofilms than healthy site-derived biofilms for each metric, which were significantly different despite overlapping confidence intervals (Fig. 2A). Further substantiation of this observation was seen in rarefaction curves using the Chao1 and Shannon metrics (Additional file 2: Figure S1A). Diversity metrics were significantly different between healthy and disease site-derived biofilms at 20,000 reads, (Phylogenetic distance p ≤ 0.05; Observed species p ≤ 0.001, Shannon p ≤ 0.001) with the exception of Chao1 (Fig. 2B, Additional file 2: Figure S1B). Outliers in the boxplots for each diversity metric are not consistently the same samples, nor are they all early or late generation samples. As diversity in generation 1 is higher than the remaining generations and may be responsible for the significant difference between healthy and disease site-derived biofilms, we repeated our alpha diversity analyses on tables that excluded generation 1 biofilms. We observed that the average alpha diversity scores of healthy and disease site-derived biofilms remain significantly different by observed species and Shannon diversity metrics but not by Faith’s phylogenetic distance (Figure S2).

**Fig. 2.**
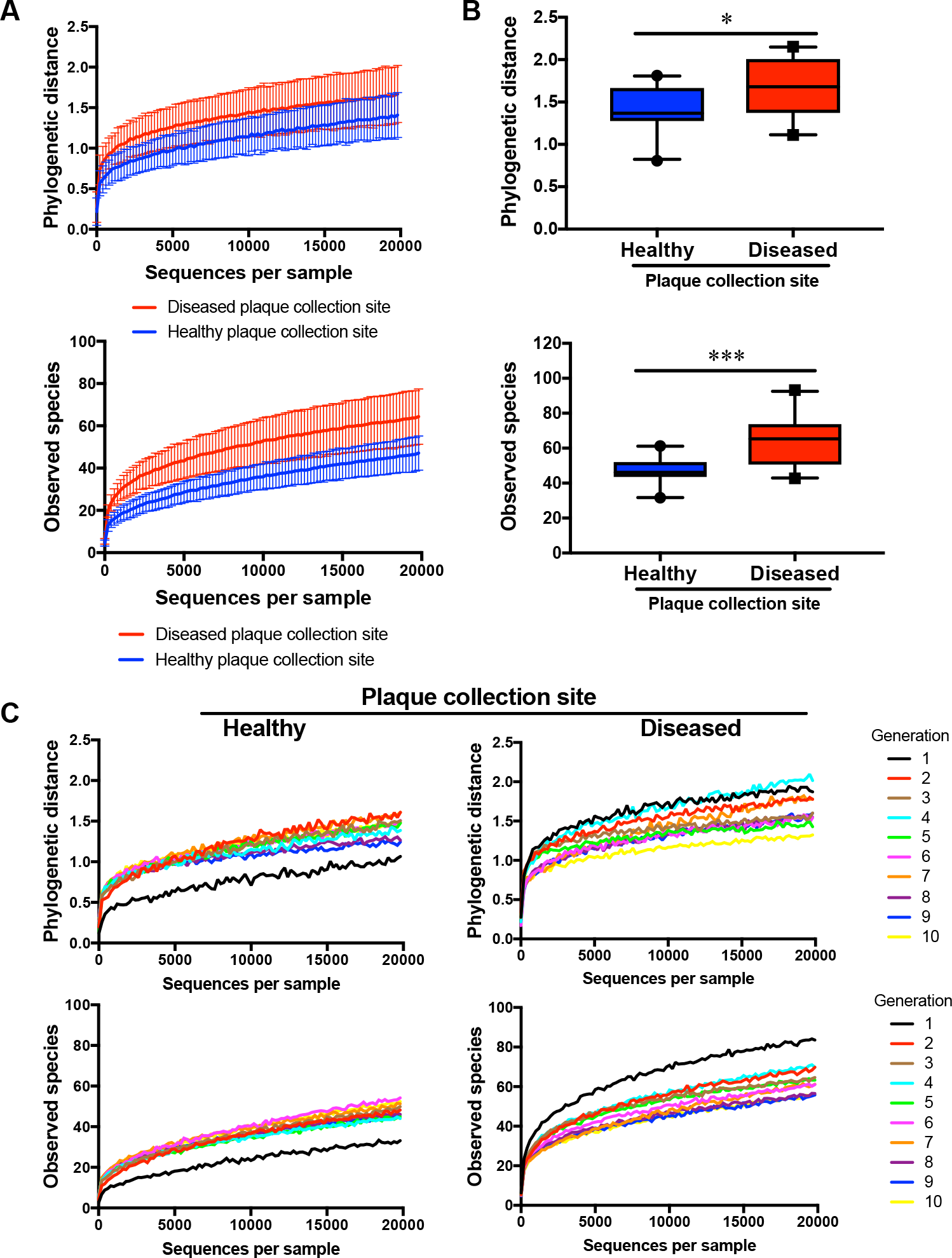
Alpha diversity of healthy and disease site-derived biofilms. (**a**) Rarefaction curves of phylogenetic distance (top) and observed species (bottom) of all ten generations combined for healthy and disease site-derived biofilms. (**b**) Phylogenetic distance (top) and observed species (bottom) are significantly different between healthy and disease site-derived biofilms at 20000 reads. (**c**) Rarefaction curves of phylogenetic distance (top) and observed species (bottom) of healthy (left) and disease (right) site-derived biofilms at each generation. Box plots show 5-95 percentile. All error bars indicate ± SD.

To see whether diversity changes substantially through repeated dispersal and reinoculation of the biofilms, we averaged the alpha diversity scores of both healthy site-derived biofilms and both disease site-derived biofilms, and plotted rarefaction curves for individual generations of healthy and disease-site derived biofilms (Fig. 2C, Additional file 2: Figure S1C). These curves reveal that for each diversity metric, between the first (black line) and second (red line) generations, diversity increases slightly in healthy site-derived biofilms yet decreases slightly in disease site-derived biofilms, while there is no substantial change in diversity between generations after the second generation in either healthy or disease site-derived biofilms. Additionally, we compared the alpha diversity scores between healthy and disease-site derived biofilms for each generation at 20,000 reads to understand if the overall significant difference in diversity is manifested at each generation (Fig. 3A,C; Additional file 3: Figure S3A,C). There were no significant differences in diversity score between healthy and disease site-derived biofilms for any generation at 20000 sequences/sample (Fig. 3B,D; Additional file 3: Figure S3B,D), but having only two samples in each group is likely insufficient power to detect statistical differences.

**Fig. 3.**
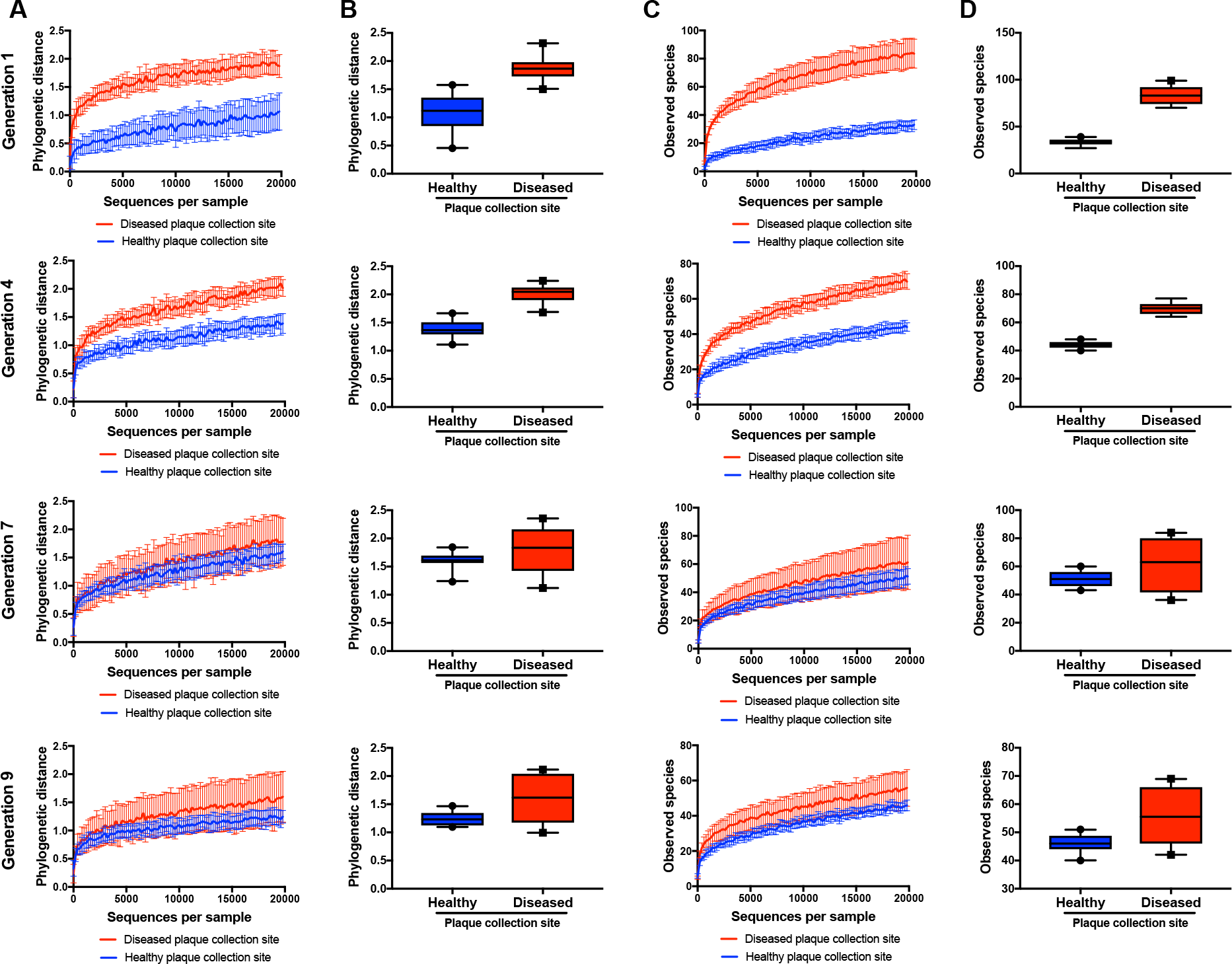
Alpha diversity of healthy and disease site-derived biofilms throughout the generations. (**a**) Phylogenetic distance rarefaction curves at generations 1, 4, 7, and 9. (**b**) Phylogenetic distance of healthy and disease site-derived biofilms by generation at 20000 reads. (**c**) Observed species rarefaction curves at generations 1, 4, 7, and 9. (**d**) Observed species in biofilms by generation at 20000 reads. N=2 biofilms per group. Box plots show 5-95 percentile. All error bars indicate ± SD.

The consistency of biofilm community composition between generations was tested by Spearman’s correlation for each lineage. The correlation coefficients demonstrated significant positive and strong correlation between generations (correlation coefficient 0.5 0. 7, p < 0.001), which was strongly supported by statistical tests (Table 1). Because low-abundance taxa were inconsistently detected between generations, we repeated the tests using a species table containing only species present at >0.1% abundance (Table 1). After removing low abundance species, the correlation coefficients between generation rose in each lineage (0.6-0. 9, p < 0.001), except between the first and tenth generations, however, still maintaining significant correlations, as observed before.

**Table 1.**
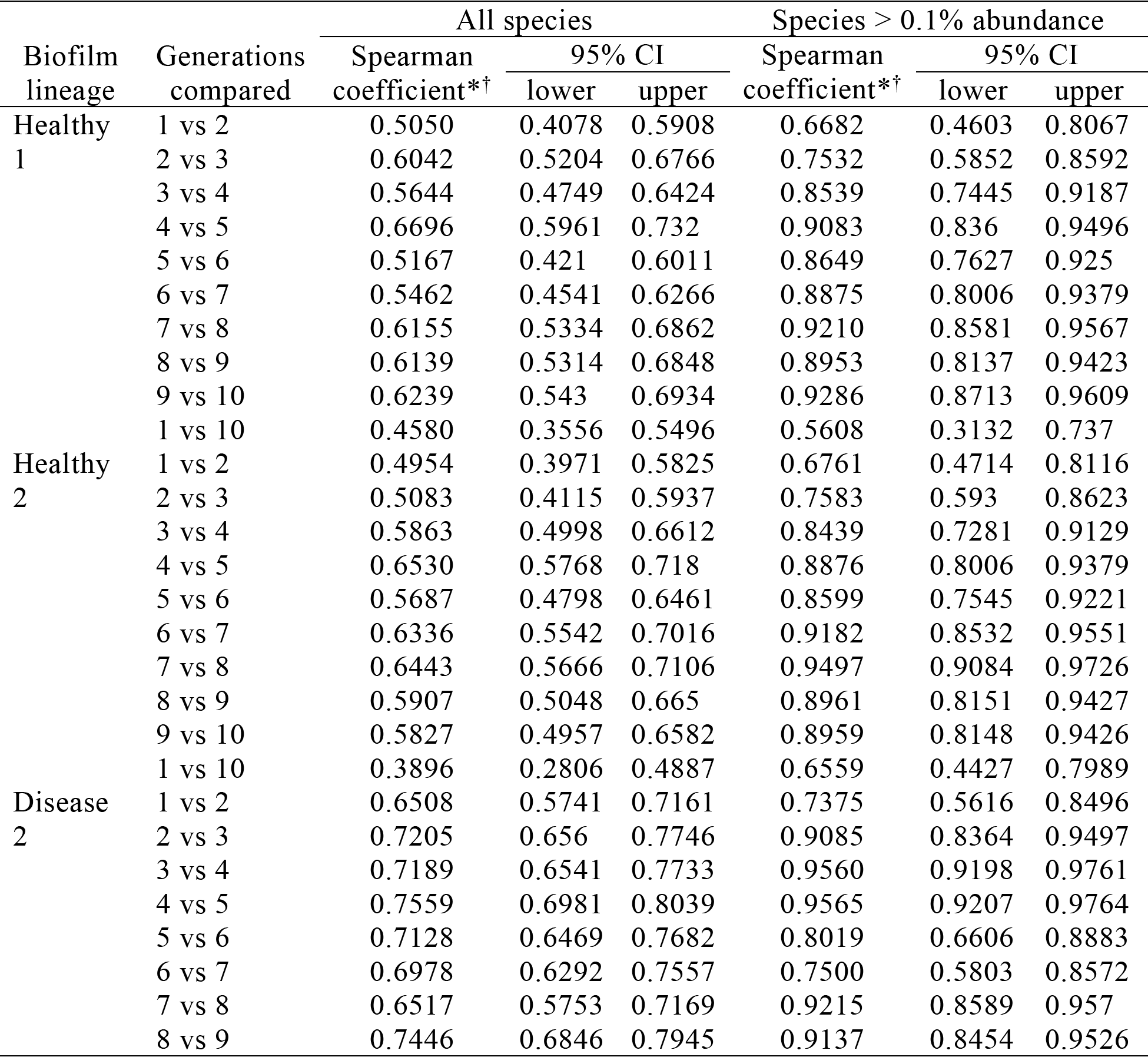
Spearman correlation of biofilm species composition between generations in each lineage.

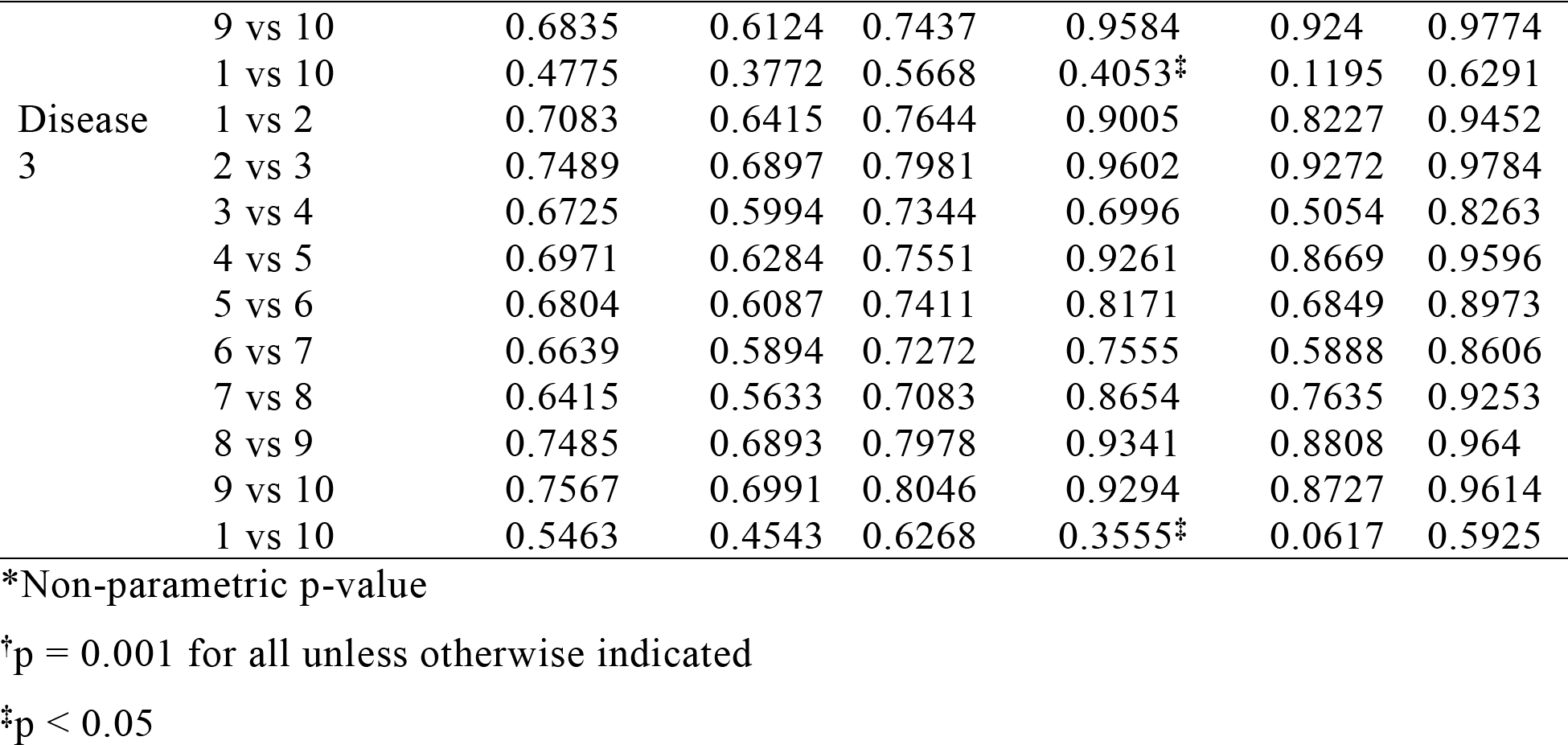

### Community diversity between healthy and diseased site-derived biofilm lineages

We also assessed community diversity differences between LAP site-and healthy site-derived biofilms (beta-diversity) to determine if the biofilms maintained a distinct community structure throughout growth in identical media. Principal coordinates analysis was used to determine separation of healthy and disease site-derived biofilms based on the weighted UniFrac metric. A clear separation of healthy and disease site-derived biofilms is seen based on the weighted UniFrac distance (Fig. 4), and the separation is significant as determined by adonis (p ≤ 0.001), indicating that there is significant community diversity between the two groups. Additionally, the Sorensen metric separated healthy and disease site-derived biofilms (p ≤ 0.001) as well as the two LAP site-derived biofilm lineages (Additional file 4: Figure S4), which suggests the initial plaque inocula from the two distinct disease patients had different microbial communities, which in fact remained distinct despite culture in the same media, while the initial healthy site plaque inocula from individual donors contained similar communities.

**Fig. 4.**
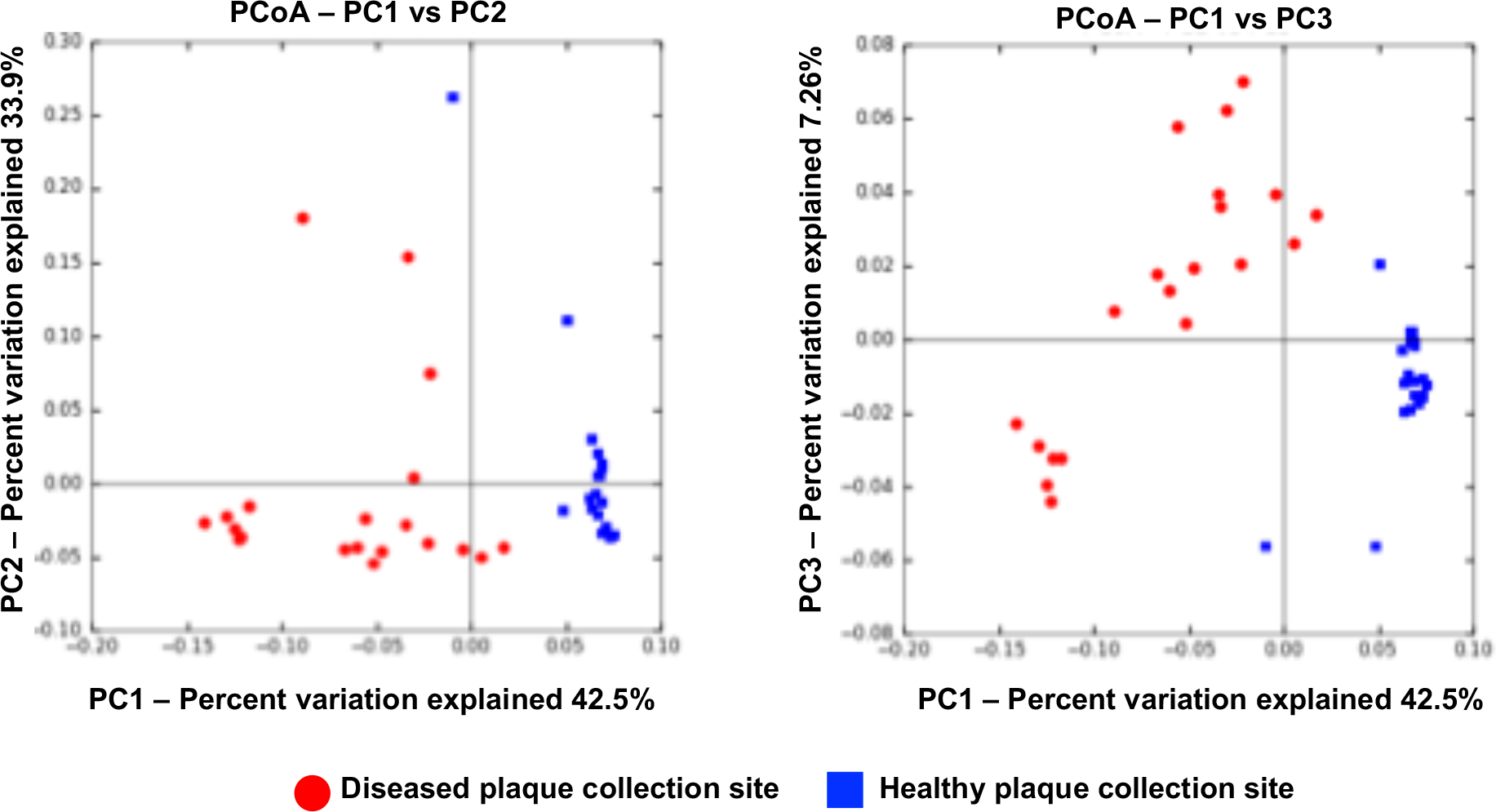
Principal coordinates analysis of beta diversity by weighted UniFrac distance of healthy and disease site-derived biofilms. Biofilms cluster based on disease status of original plaque inocula, and group separation is significant based on adonis analysis (p < 0.001).

To test the strength of these results we also performed jack-knifed beta diversity using weighted UniFrac and Sorensen metrics, and performed PCoA on the matrices (Additional file 5: Figure S5A,B), with results very similar to that in Figures 3 and S4. This repeated calculation of beta-diversity by leaving out a randomly selected sample each time reveals if there are samples that strongly affect community diversity, and may therefore be outliers in the dataset. Repeated analyses of the weighted UniFrac metric had very little variation, with only six samples with ellipsoids demonstrating plot location variability, strongly supporting the separation between healthy and disease site-derived biofilms. In contrast, the Sorensen metric had substantial variation within the healthy biofilms and within the disease site biofilms, as the ellipsoids extend predominantly along the PC3 axis, which separates the generations of each group, rather than along the PC1 axis, which separates healthy and LAP-derived biofilms. For both diversity metrics, however, consensus UPGMA trees group the healthy biofilms separate from the disease site-derived biofilms, although biofilms are not distributed in the trees here by chronological order (Additional file 6: Figure S6). These analyses support the observation that distinct community composition between healthy and LAP site-derived biofilms are maintained through the generations.

### Differentially abundant taxa between healthy and disease site-derived biofilms

Methods to determine species significantly different between different environments can be particularly sensitive to low-abundance taxa. Therefore, we determined differentially abundant taxa between healthy and disease-site derived biofilms by comparing the difference in species mean proportions using the program STAMP, first by comparing all detected species (Fig. 5) and then comparing only species that were detected at > 0.1% abundance (Additional file 7: Figure S7). Removing organisms present in less than 0.1% abundance resulted in a species table with 43 species, just 17% of the 257 total species detected, however this had no effect on the species that met cut-off criteria for differential abundance (q ≤ 0.05 and an effect size > 2). Fourteen species were differentially abundant between healthy and disease site-derived biofilms (Figures 5, S7). Seven species were more abundant in healthy site-derived biofilms (blue bars/dots), while six were more abundant in disease site-derived biofilms (red bars/dots).

**Fig. 5.**
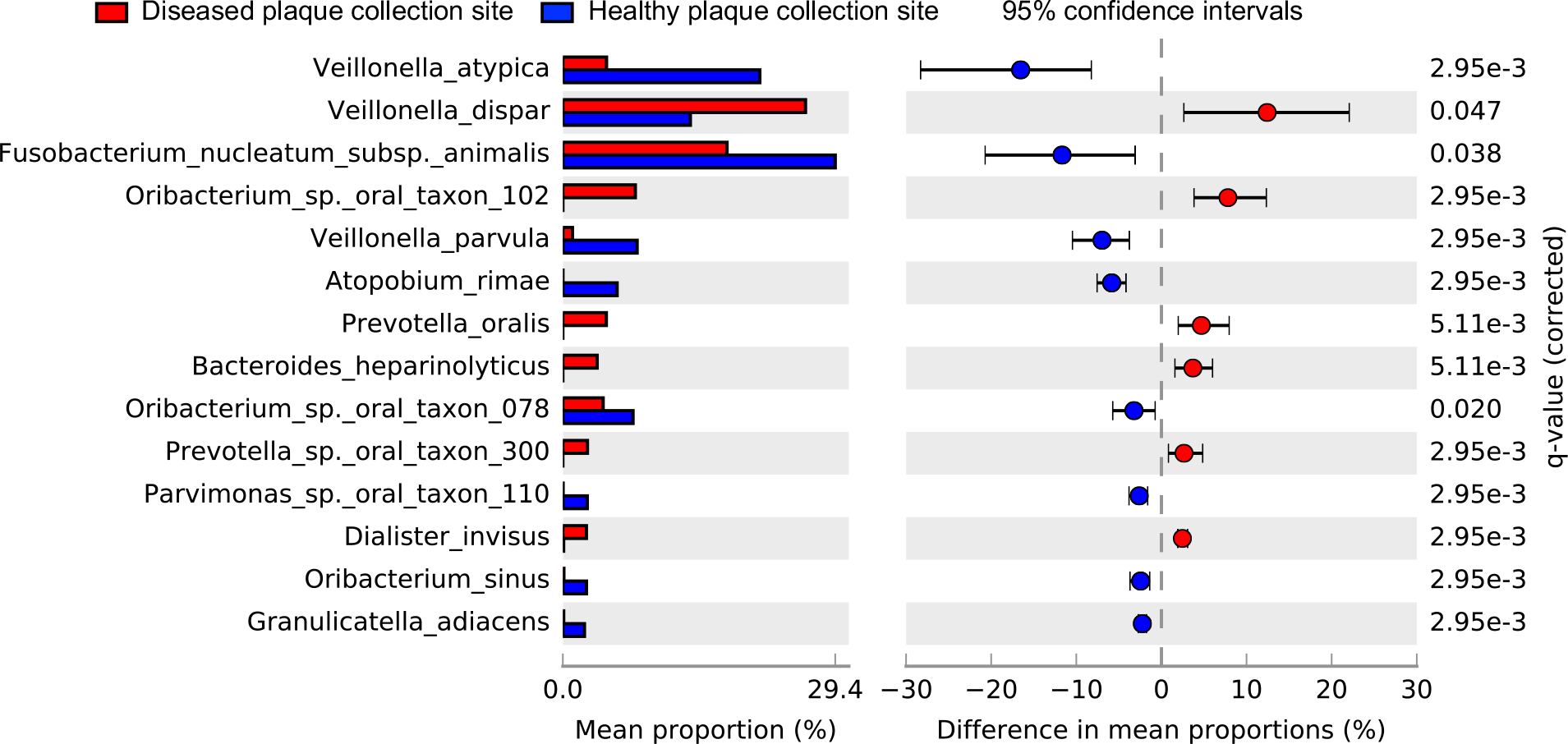
Differential abundance of species between healthy and disease site-derived biofilms. Species statistically different (q ≤ 0.05) and with an effect size (DP) >1 by analysis in STAMP are considered differentially abundant.

Comparing healthy and disease site-derived biofilms of each generation revealed a similar profile difference to that of the combined generations. Species that were significantly differentially abundant in at least seven generations included *Atopobium rimae, Fusobacterium nucleatum* subsp. *animalis, Veillonella atypica* and *V. parvula* in healthy site-derived biofilms, and *Oribacterium* sp. oral taxon 102, *Prevotella oralis*, and *Veillonella dispar* in disease site-derived biofilms. Of note, *V. dispar* was significantly more abundant in healthy site-derived biofilms for the first two generations, then for the remaining 8 generations was more abundant in disease-site derived biofilms, while *F. nucleatum* subsp. *animalis* displayed opposite dynamics, initially more abundant in disease-site derived biofilms, but after the first generation was more abundant in healthy site-derived biofilms.

### Core communities in healthy and disease site-derived biofilms

We determined the species that were detected in every generation of both healthy plaque-derived lineages to define the “healthy core community”, and the species detected in every generation of both disease plaque-derived lineages to define the “disease core community”. Healthy site-derived biofilms had eleven core species: *Fusobacterium nucleatum* subsp. *animalis, Fusobacterium* sp. oral taxon 204, *Granulicatella adiacens, Lachnoanaerobaculum saburreum, Oribacterium* sp. oral taxon 078, *Prevotella melaninogenica, Rothia mucilaginosa, Streptococcus sanguinis, Veillonella atypica, Veillonella dispar*, and *Veillonella parvula*, while disease site-derived biofilms had fourteen core species: *Atopobium parvulua, Campylobacter curvus, Capnocytophaga* sp. oral taxon 338, *Dialister invisus, Fusobacterium nucleatum* subsp. *animalis*, *Fusobacterium nucleatum* subsp. *nucleatum, Fusobacterium periodonticum, Fusobacterium* sp. oral taxon 204, *Granulicatella adiacens, Lachnoanaerobaculum saburreum, Oribacterium* sp. oral taxon 078, *Veillonella atypica, Veillonella dispar*, and *Veillonella parvula.* Several species included in one but not both core communities are missing in a single generation in one or both of the lineages, so may actually be present and undetected.

### Propagation of uncultured taxa

A unique advantage of growing complex plaque-derived biofilms compared to defined polyspecies biofilms is the inclusion of rare and uncultured microbes. The role of these organisms in oral biofilm structure and activity is difficult to study *in vitro* due to the difficulty in culturing these organisms in the lab, but next generation sequencing has opened a way to study their abundance, distribution, and gene content *in silico*, which may facilitate their independent culture in a laboratory setting. Both the LAP site-derived and healthy site-derived biofilms contained numerous uncultured oral organisms identified in the Human Oral Microbiome Database (HOMD). Several uncultured species were detected in biofilms throughout the generations, including *Peptostreptococcaceae* [XI][G-3] sp. oral taxon 495 *Prevotella* sp. oral taxon 300, *Prevotella* sp. oral taxon 304, Prevotella sp. oral taxon 315, *Fusobacterium* sp. oral taxon 204, *Fusobacterium* sp. oral taxon 205, *Stomatobaculum* sp. oral taxon 097, and *Stomatobaculum* sp. oral taxon 910. Members of the phylum TM7 (now *Candidatus Saccharibacteria*), which has only one cultured member **[46]**, were detected in both healthy site-derived biofilms, up to 7 generations and up to 10 generations, respectively, while they were detected in disease site-derived biofilms up to 4 and 5 generations, respectively. Additionally, members of the uncultured phylum SR1 were detected in the 4^th^ generation of one healthy site-derived biofilm, and in the 3^rd^, 7^th^ and 9^th^ generation of one disease site-derived biofilm. This indicates that our biofilm model is capable of sustaining growth of these poorly characterized organisms and may be useful for their *in vitro* study.

## Discussion

With the emphasis growing on how oral microbial community interactions influence health and disease, *in vitro* models of complex oral biofilms are likely to become increasingly used. Such models need to maintain viable communities for long-term studies and reproduce consistent communities to reduce variability in study conditions. The developers of the model we used here determined that the biofilms could not be reliably grown longer than ten days, because the biofilms would release from the disks [28], necessitating the development of a method to perpetuate the biofilms [33]. Here we have demonstrated longevity and reproducibility of biofilms seeded from a single source by Next Generation Sequencing technology, and shown that stationary growth of subgingival plaque-derived biofilms can maintain complex multi-species microbial communities including uncultivatable species, even sustaining signatures of health and disease when grown in identical media. These results are highly promising for use of this model as a complex oral biofilm for studies of plaque development, microbial-microbial interactions, and microbial-host interactions, while further development and refinement of the model are likely to broaden and enhance its applications.

Long-term *in vitro* biofilm growth models are particularly appealing for studies using patient samples or animal models, which may last weeks to months. In these circumstances, it may not be possible to recall the same plaque donor, and patients with periodontal disease who undergo treatment cannot be re-sampled. We recently published an application of this model to stimulating patient peripheral blood mononuclear cells [35], where patients fulfilling inclusion criteria were enrolled over several months. We aimed to measure immune cell responses to stimulation with biofilms from the same source to reduce potential variability in responses, in contrast to our study that used fresh plaque collected from a new patient to stimulate each new set of patient samples [17]. Maintaining several biofilm lineages allowed us to have biofilms grown from the same source ready for stimulation each time patient blood samples were collected. Additionally, serially grown biofilms can be used for murine oral infection models of periodontal disease, which require serial infection over 3-24 weeks [15,16,47] to develop symptoms. Multi-species infection models could use serially grown biofilms from human plaque, rather than independently growing several bacterial species and combining them just before infection, which is the current standard [15,16].

While comparison of all four biofilm lineages to their initial plaque inocula would be beneficial for understanding what proportion of the original diversity is established in culture, we did not have any remaining inocula from 3 of our samples to sequence. However, a comparison with initial plaque inocula was previously done for our model [28,33], and our goal was instead to demonstrate consistent and long-term reproducible biofilm growth. There are no *in vitro* oral biofilm growth models that support greater than 80% of the species present in the initial plaque or saliva inoculum, and based on assessments by Walker and Sedlacek [28] and Shaddox, *et al.* [33] as well as comparison of the plaque sample used to inoculate DB3 with the biofilms of that lineage here, our model our model supports 53-80% of the initial community. This is comparable to four independent oral biofilm microcosm models that reproduced 64% [29], 78% [30], 62% [31] and 44% [32] of the OTUs present in the initial inocula. Biofilms grown from plaque rather than saliva appear to lose a greater amount of diversity, a phenomenon that suggests that there are some fundamental components necessary for *in vivo* plaque growth that have not yet been elucidated and replicated *in vitro*. For example, Fernandez y Mostajo, *et al.* [32] found that supplementing growth media with high amounts of serum was necessary to support growth of *Treponema* species, and further studies of plaque biofilm ecology are clearly needed to improve *in vitro* models. Despite this limitation, we were able to demonstrate here that serially-grown biofilms could consistently reproduce similar, stable communities, through several generations.

It was recently shown in *E. coli* that cell lineage history was a stronger predictor of overall cell metabolic activity than spatial proximity [48]. If this holds true for other bacterial species, especially when growing in complex biofilm communities such as these, selecting the source of biofilm inoculum becomes important, and thus, plaque biofilm inoculum may be preferential to saliva inoculum or defined species inoculum for seeding *in vitro* biofilms. The bacteria in dental plaque are already established, transcriptionally and translationally, for biofilm growth and are likely to be more closely related to each other than are bacterial cells in saliva because of constant mixing and turnover of saliva. Further, initial characterization of the subgingival plaque model used in our study demonstrated that biofilm growth was inhibited when sterilized saliva from individuals other than the dental plaque donor were used to initiate biofilm growth [28]. This suggests that an individual’s microbiota are optimally adapted for surface attachment in their native saliva, while salivary components from other donors may contain inhibitory molecules such as antibodies or small antimicrobial peptides that the microbiota are not adapted to combat. Therefore, dental plaque inocula may be more likely develop *in vitro* biofilms that maintain metabolic characteristics of healthy and disease site plaque, and thus, this warrants further investigation.

Although it is uncommon for *in vitro* oral biofilm studies to report biofilm community composition beyond 72 hours of growth [29,31], allowing the biofilms to grow to maturity (seven to ten days for this model [33]) may be critical to maintaining community profiles for propagating the biofilms, as longer growth periods allow slow-growing, low-abundance and “late colonizer” species to reach sufficient levels that they are not out-competed by rapid-growing species, or lost by chance from the small aliquots used to inoculate subsequent generations. Saliva-and plaque-derived biofilms as well as subgingival plaque develop continuously over at least seven days [28,30,49], which supports using more mature biofilms (older than 72 hours) in studies of biofilm-host interactions, as older biofilms are more representative of plaque collected from subgingival sites, which themselves require days to develop familiar community profiles.

While community changes between generations due to inconsistent detection of low abundance species may appear to make the system unreliable, it is similar to the recently reported dynamics of oral microbial communities [50,51]. Hall, *et al.* [50] demonstrated that a core set of microbes were consistently present over time in temporally sampled supragingival plaque, tongue swabs, and saliva, but that there was substantial change in low abundance organisms, which caused community drift. Our biofilm generations similarly have a core community with a few species that comprise a majority of the reads in each sample, and the changes in the community are predominantly due to the low-abundance species. Additionally, our biofilm communities do not appear to drift beyond a defined profile, just as is reported in plaque communities over time. These dynamics suggest that oral biofilms, while influenced by their environment, are stable and resilient communities, wherein the metabolic and physical interactions of the members prevent drastic community profile changes.

The microbiological composition of LAP site and healthy site plaque has been characterized in this population [52], and we compared the composition of our biofilm samples to this to see how well the biofilms reflect the known characteristics of LAP site and healthy site plaque. Many of the species that were significantly more abundant in LAP plaque samples were fastidious asaccharolytic organisms, which propagate poorly in our model. Even though we do not detect high levels of species traditionally associated with periodontitis in our disease site-derived biofilm samples, we do detect several species that were recently associated with LAP by NGS [52], including *Solobacterium moorei, Parvimonas micra, Capnocytophaga* sp., *Filifactor alocis, Aggregatibacter actinomycetemcomitans* and *Tannerella forsythia.* There is very little overlap between the core species in the biofilm lineages and the most abundant species reported in LAP plaque and healthy control plaque samples using HOMIM technology [52], despite consistency between HOMIM and HOMINGS technologies [36]. However, there is substantial overlap between the core communities of healthy and periodontitis sites as has been reported previously [5].

Indeed, beta-diversity analysis of the plaque samples discussed in Shaddox, *et al.* [17], reveal significant differences between LAP site plaque and healthy patient healthy site plaque (p < 0.05 by Bray-Curtis similarity, Sorensen-Dice index and Jaccard index by adonis test). Although it is not possible to directly compare HOMIM and HOMINGS data, the significant difference within each dataset supports that plaque at healthy and disease-affected sites harbor uniquely diverse communities, which has also been reported in other *in vitro* biofilm models [32]. An unrelated study of dental plaque communities in periodontal health and disease found that periodontitis-associated communities cluster into two groups related to periodontitis clinical parameters [53], while another found several distinct communities within generalized aggressive periodontitis site plaque [18]. Distinct microbial profiles within LAP plaque samples have not yet been investigated, but is a possible explanation for the differences we observe.

*Fusobacterium nucleatum* subsp. *animalis* and *Veillonella dispar* were the only two species that started highly abundant in biofilms from one source and during culture decreased in abundance in those lineages and became highly abundant in the other. Both species are common in sites of disease and health [5], and may have flexible metabolic requirements that allow them to rapidly multiply in the presence of a wide variety of other bacterial species. Promisingly, the healthy and diseased site-derived biofilms still maintain a differentiated profile through the generations, and a potential unique metabolic profile as well, evidenced by the distinct difference in smell, although this was not directly tested here. This may be related to the production of short chain volatile fatty acids and volatile sulfur compounds, which are attributed to proteolytic, disease-associated species [54]. Recent reports on the transcriptional activity of biofilm plaque from healthy and periodontitis-affected sites suggest that community activity is more uniform across disease sites than is microbial profile [6,7,55], and further investigation of protein expression and metabolic output in healthy and disease-site derived biofilms is warranted [56].

In conclusion, we have demonstrated that our subgingival oral biofilm growth and perpetuation model can maintain reliable communities and differentiated health/disease microbial profiles for at least 80 days, and mimics dynamics of natural plaque. This method of biofilm perpetuation may be useful for growing biofilms to use in *in vivo* oral infection studies, patient immune cell stimulation studies, biofilm transcriptional, translational, and metabolic studies, and to study the effects of anti-biofilm agents.

## Declarations

### Ethics approval and consent to participate

Dental plaque samples used in this study were collected as part of a larger clinical study approved by the University of Florida Institutional Review Board (IRB#201400349) and registered at clinicaltrials.gov (NCT01330719). All participants/LAR signed an informed consent form.

### Consent for publication

Not applicable

### Availability of data and material

All data generated or analyzed during this study are included in this published article and its supplementary information files. Raw data is available upon request.

### Competing interests

The authors declare that they have no competing interests

### Funding

This work was supported by NIH NIDCR R01DE019456 (LMS) and NIH NIDCR T90 DE021990 (IMV). The funding body had no role in the design of the study and collection, analysis, and interpretation of data or in writing the manuscript.

### Authors’ contributions

LMS and IMV designed the study. IMV performed biofilm growth experiments, bioinformatics analyses and drafted manuscript. LMS performed diagnosis and sample collection of subjects, oversaw study and revised manuscript. All authors read and approved the final manuscript.

## Acknowledgements

We thank Dr. Ann Progulske-Fox for the use of her anaerobic chamber during this study.

## Additional files

**Additional File 1: Table S1**. Per sample total read counts and absolute read counts for each species and genus detected (species table from HOMINGS analysis).

**Additional File 2: Table S2**. Per sample total read counts and absolute read counts for each species and genus detected by QIIME analysis of raw fasta files. Tab 1 (otu_table_L6) is all taxonomic assignments down to genus level. Tab 2 (otu_table_L7) is all taxonomic assignments down to species level. Tab 3 (CompHOMINGS-qiime) has total, matched, and unmatched read counts for each biofilm for HOMINGS and QIIME analysis, as well as the total number of species detected by each method.

**Additional File 3: Figure S1.**
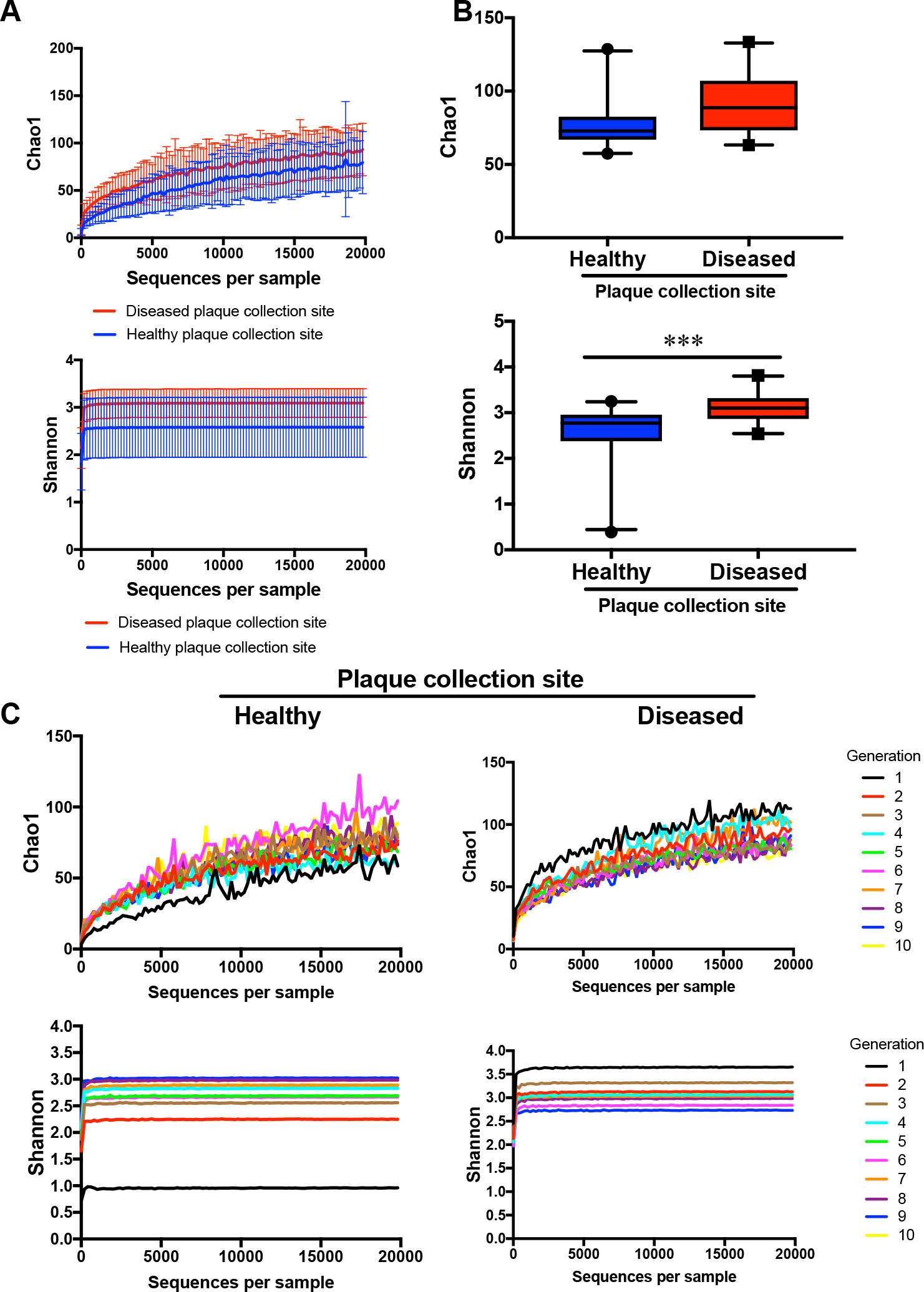
Alpha diversity of healthy and disease site-derived biofilms. (**a**) Rarefaction curves of Chao1 index (top) and Shannon index (bottom) of all combined generations for healthy and disease site-derived biofilms. (**b**) Chao1 index (top) and Shannon index (bottom) are significantly different between healthy and disease site-derived biofilms at 20000 reads. (**c**) Rarefaction curves of Chao1 index (top) and Shannon index (bottom) of healthy (left) and disease (right) site-derived biofilms at each generation. Box plots show 5-95 percentile. All error bars indicate ± SD.

**Additional File 4: Figure S2.**
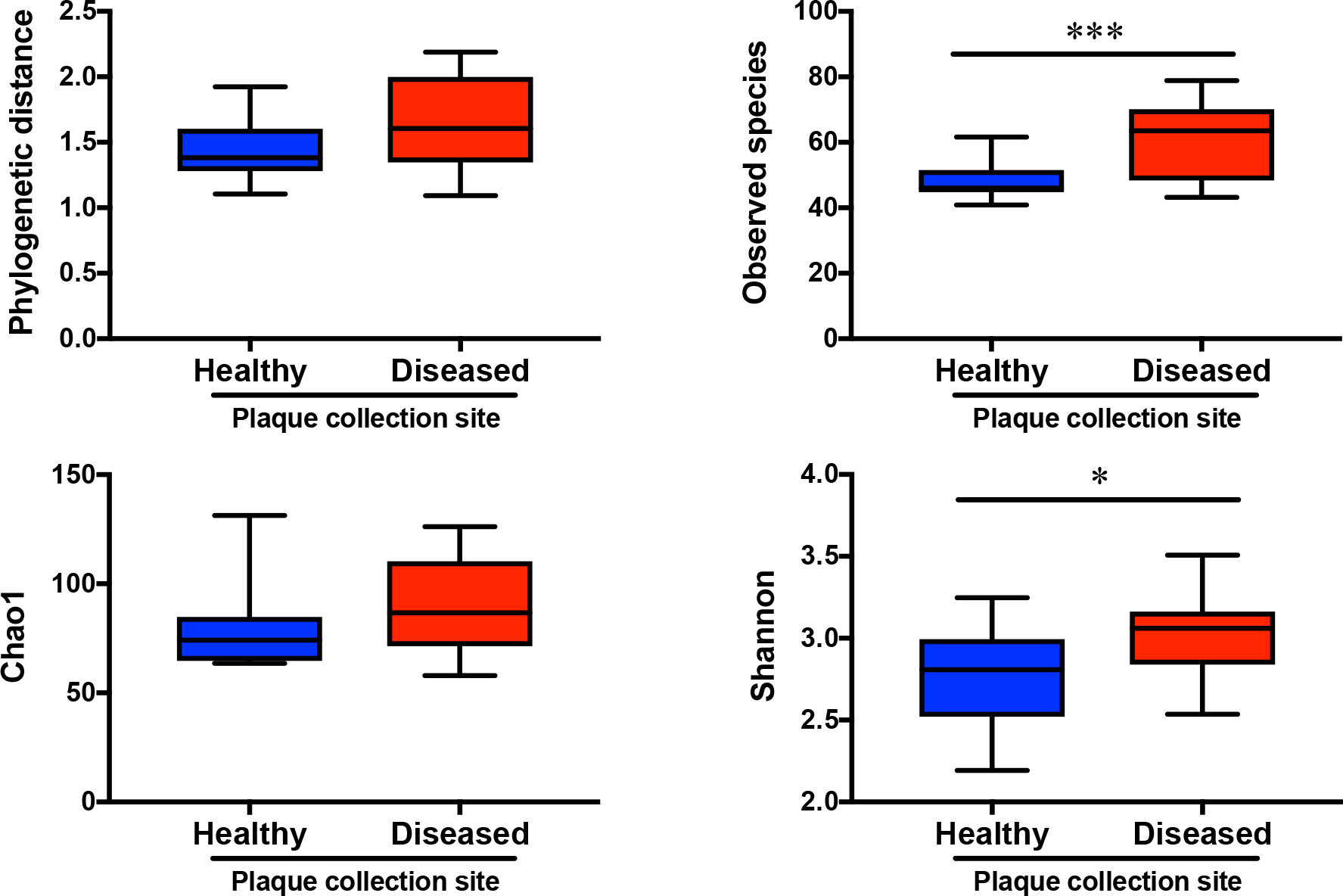
Alpha diversity of healthy and disease-site derived biofilms generations 2-10. Alpha diversity score for Faith’s phylogenetic distance, Observed species, Chao1 and Shannon indexes at 20000 reads averaged across generations 2-10. Box plots show 5-95 percentile. All error bars indicate ± SD. * p < 0.05, *** p < 0.001.

**Additional File 5: Figure S3.**
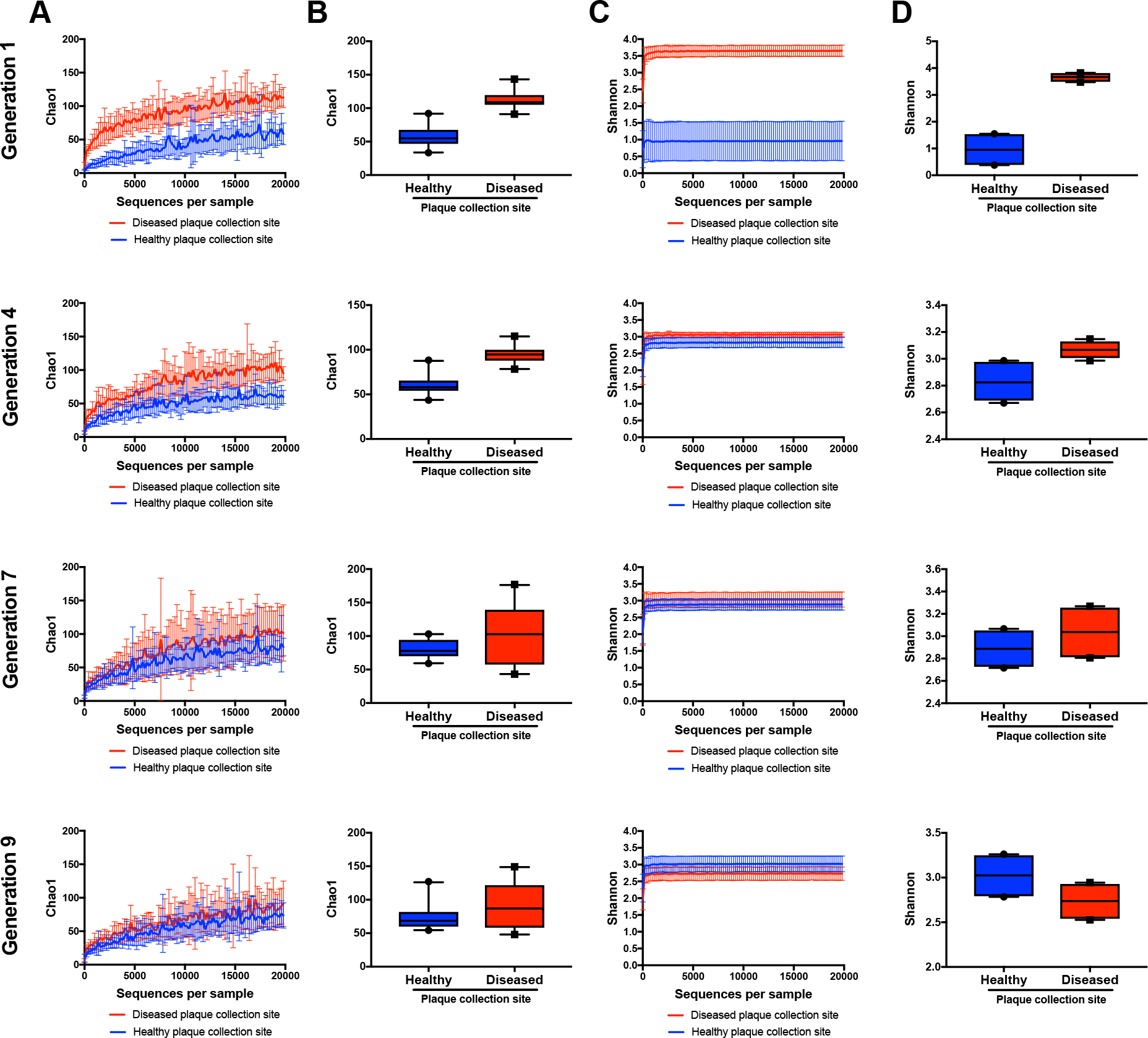
Alpha diversity of healthy and disease site-derived biofilms throughout the generations. (**a**) Chao1 index rarefaction curves at generations 1, 4, 7, and 9. (**b**) Chao1 index of biofilms by generation at 20000 reads. (**c**) Shannon index rarefaction curves at generations 1, 4, 7, and 9. (**d**) Shannon index in biofilms by generation at 20000 reads. N=2 biofilms per group so statistical significance could not be calculated. Box plots show 5-95 percentile. All error bars indicate ± SD.

**Additional File 6: Figure S4.**
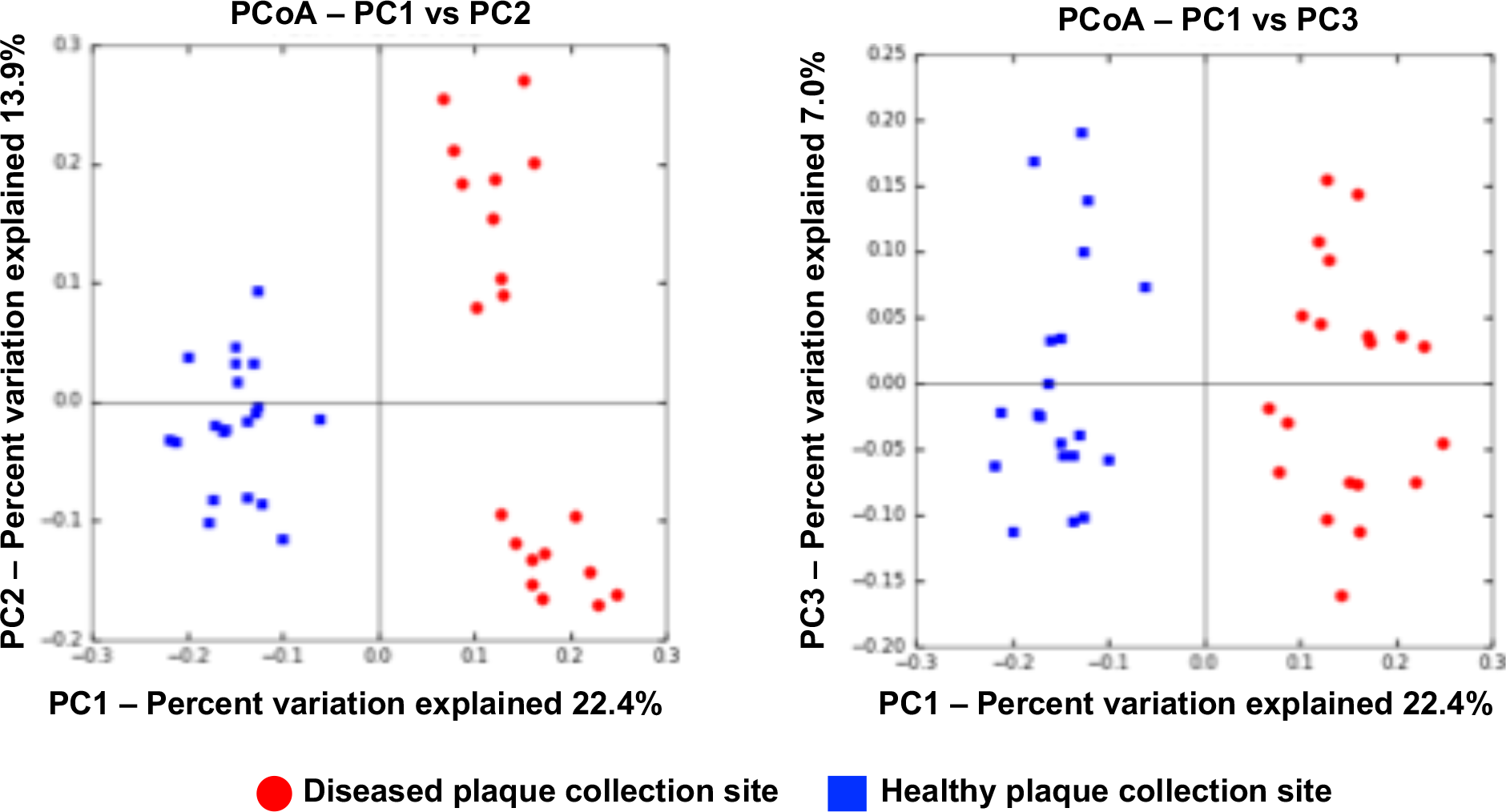
Principal coordinates analysis of beta diversity by Sorensen distance of healthy and disease site-derived biofilms. Biofilms cluster based on disease status of original plaque inocula and group separation is significant based on adonis analysis (p < 0.001). Additionally, disease site-derived biofilms cluster into 2 groups based on plaque inocula donor.

**Additional File 7: Figure S5.**
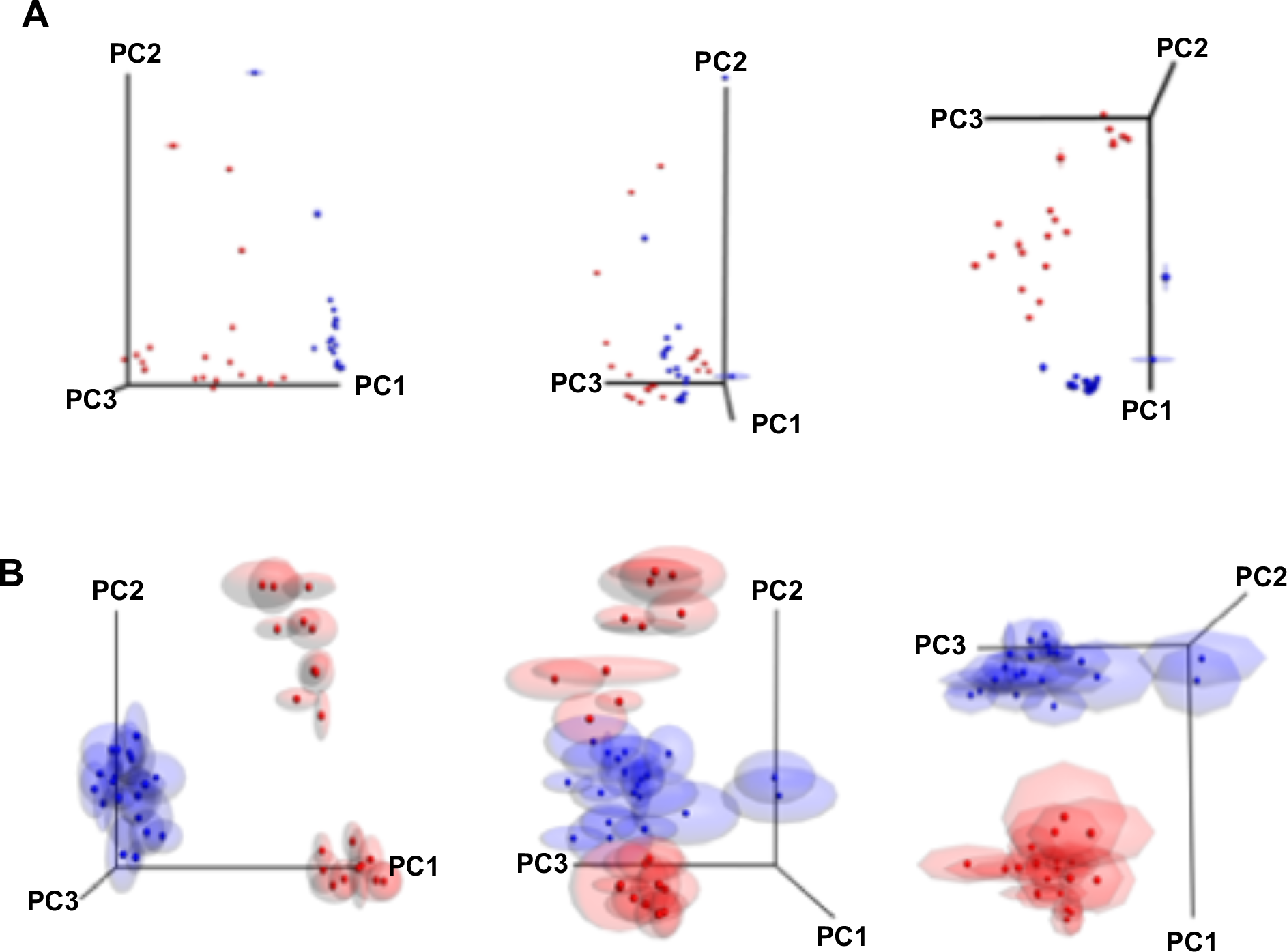
Principal coordinates analysis of jack-knifed beta diversity by weighted UniFrac and Sorensen distances of biofilms. Biofilms cluster based on disease status of original plaque inocula and disease site-derived biofilms separate based on donor. (**a**) Biofilm coordinates exhibit very little variation in PCoA based on weighted UniFrac distance. (**b**) Biofilm coordinates demonstrate variation in PCoA based on Sorensen distance but retain distinct clustering of healthy and disease site-derived groups. Additionally, disease site-derived biofilms likewise retain distinct separation into groups based on plaque inocula donor.

**Additional File 8: Figure S6.**
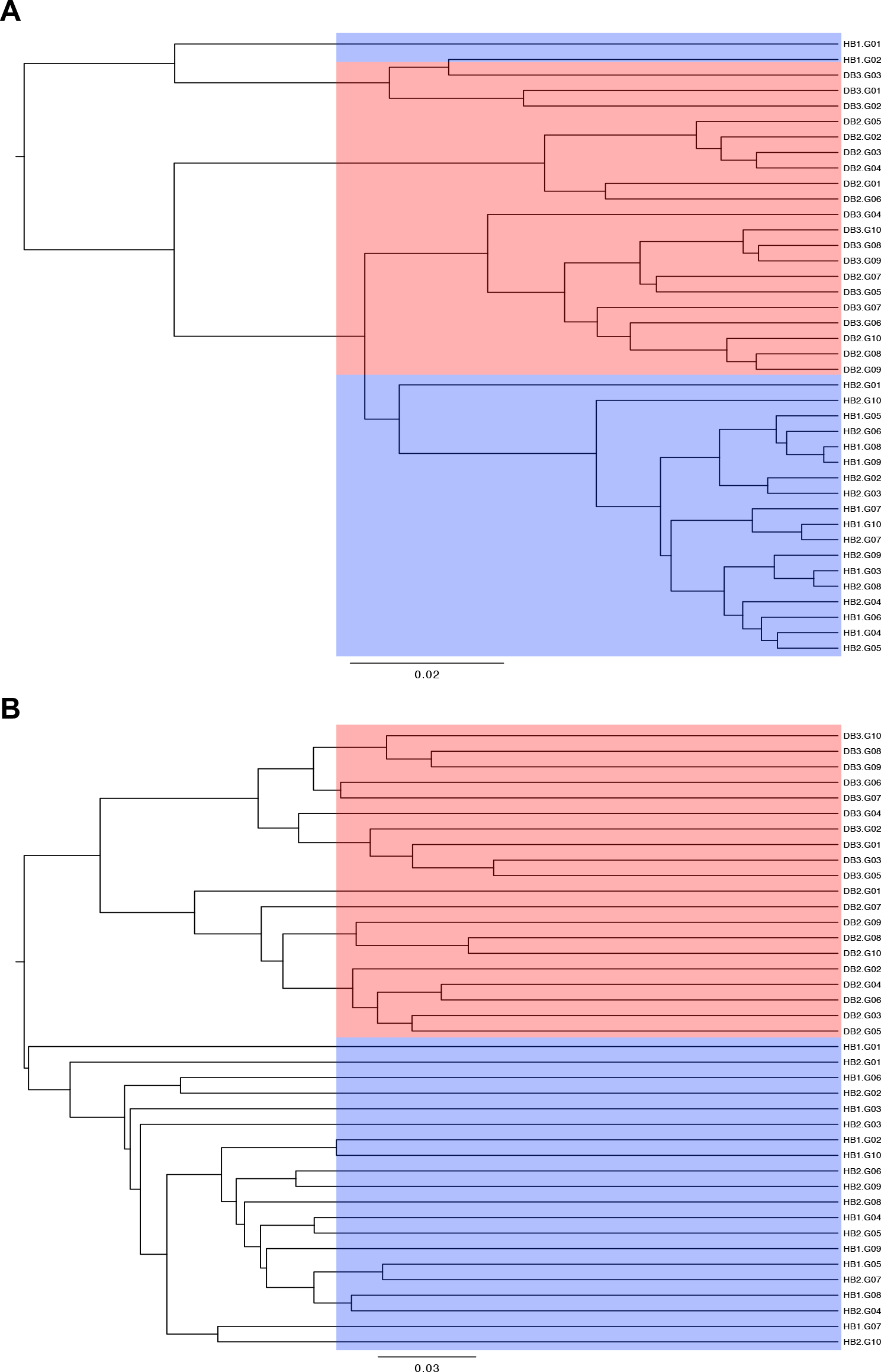
UPGMA consensus trees of jackknifed beta-diversity measurements. (**a**) Weighted UniFrac distance groups several early biofilm generations despite differences in plaque source, but otherwise distinguishes between healthy and disease site-derived biofilm lineages. (**b**) Sorensen distance clearly distinguished between healthy and disease site-derived biofilm lineages.

**Additional File 9: Figure S7.**
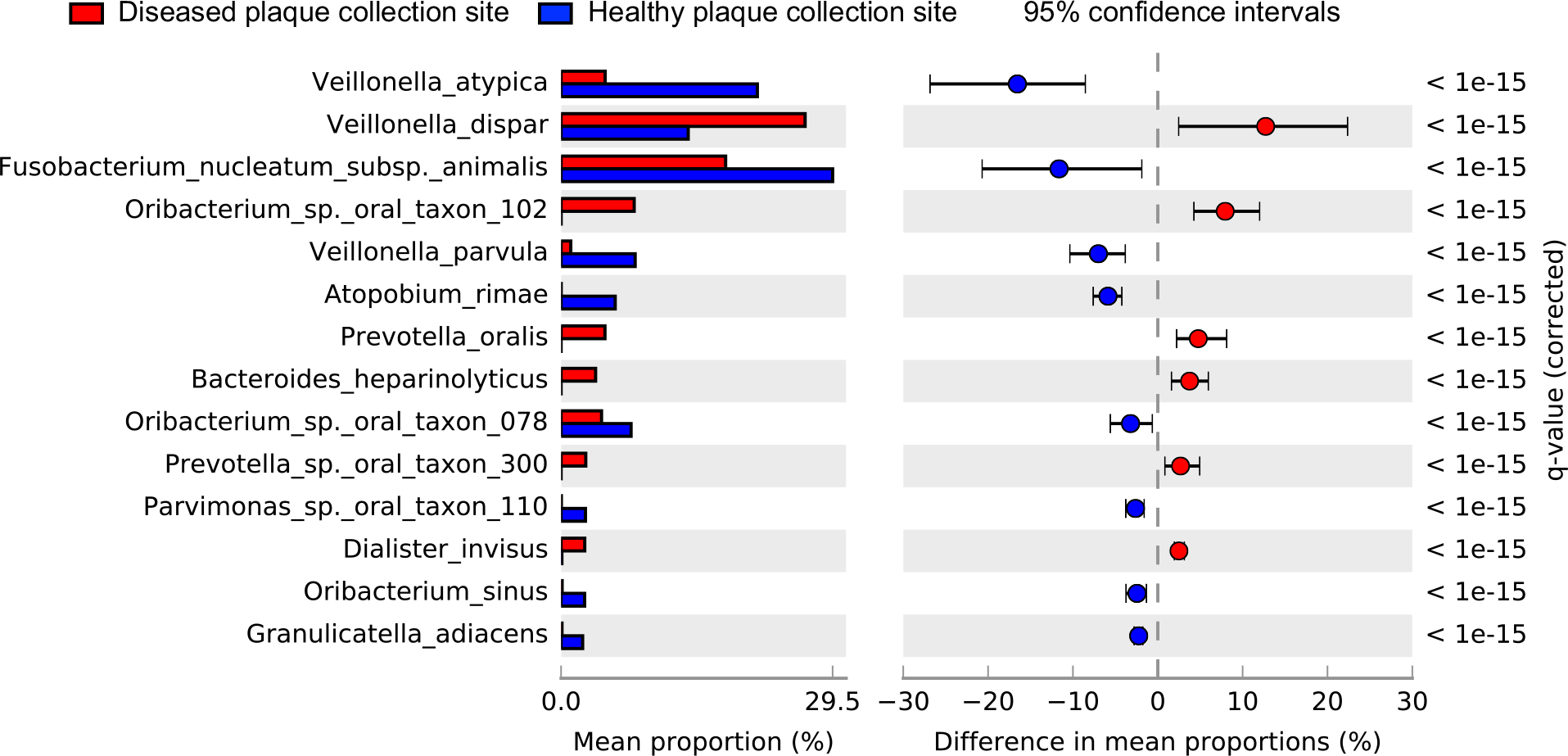
Differential abundance of species present at >0.1% abundance between healthy and disease site-derived biofilms. Species statistically different (q ≤ 0.05) and with an effect size (DP) >1 by analysis in STAMP are considered differentially abundant.

